# Functionally overlapping variants control TB susceptibility in Collaborative Cross mice

**DOI:** 10.1101/785725

**Authors:** Clare M. Smith, Megan K. Proulx, Rocky Lai, Michael C. Kiritsy, Timothy A Bell, Pablo Hock, Fernando Pardo-Manuel de Villena, Martin T. Ferris, Richard E. Baker, Samuel M. Behar, Christopher M. Sassetti

## Abstract

Host genetics plays an important role in determining the outcome of *Mycobacterium tuberculosis* (Mtb) infection. We previously found that Collaborative Cross mouse strains differ in their susceptibility to Mtb, and that the CC042/GeniUnc (CC042) strain suffered from a rapidly progressive disease and failed to produce the protective cytokine, IFN*γ*, in the lung. Here, we used parallel genetic and immunological approaches to investigate the basis of CC042 susceptibility. Using a population derived from a CC001/Unc (CC001) × CC042 intercross, we mapped four QTL underlying Tuberculosis ImmunoPhenotypes (*Tip1-4*). These included 2 major effect QTL on Chromosome 7 (*Tip1* and *Tip2*) that were associated with bacterial burden. *Tip2*, along with *Tip3* (Chromosome 15) and *Tip4* (Chromosome 16) also correlated with IFN*γ* production following infection, whereas *Tip1* appeared to control an IFN*γ*-independent mechanism of bacterial control. Further immunological characterization revealed that CC042 animals recruited relatively few antigen-specific T cells to the lung and these T cells failed to express the Integrin alpha L (α_L_; i.e., CD11a), which contributes to T cell activation and migration. These defects could be explained by a CC042 private variant in the *Itgal* gene, which encodes CD11a, and is found within the *Tip2* interval. This 15bp deletion leads to aberrant mRNA splicing and is predicted to result in a truncated protein product. The *Itgal^CC042^* genotype was associated with all measured disease traits, indicating that this variant is a major determinant of susceptibility in CC042. The combined effect of functionally distinct *Tip* variants likely explains the profound susceptibility of CC042 and highlights the multigenic nature of TB control in the Collaborative Cross.

**Importance statement:** The variable outcome of *Mycobacterium tuberculosis* infection observed natural populations is difficult to model in genetically homogenous small animal models. The newly-developed Collaborative Cross (CC) represents a reproducible panel of genetically-diverse mice that display a broad range of phenotypic responses to infection. We explored the genetic basis of this variation, focusing on a CC line that is highly susceptible to *M. tuberculosis* infection. This study identified multiple quantitative trait loci associated with bacterial control and cytokine production, including one that is caused by a novel loss-of-function mutation in the *Itgal* gene that is necessary for T cell recruitment to the infected lung. These studies verify the multigenic control of mycobacterial disease in the CC panel, identify genetic loci controlling diverse aspects of pathogenesis, and highlight the utility of the CC resource.

## Introduction

Nearly one quarter of the world’s population has been exposed to *Mycobacterium tuberculosis* (Mtb), yet less than 10% of these exposures progress to clinical disease (1). The rational design of more effective interventions requires understanding the factors that determine the outcome this interaction. A large body of evidence supports an important role for host genetics in determining disease progression, including classic twin studies (2, 3), linkage analyses (4–8), and both case-control (9, 10) and genome-wide association studies (11, 12). However, the genetic variants that determine the risk of adult pulmonary disease remain elusive due both to the complexity of factors influencing clinical outcome and the lack of model systems that reflect the diversity of natural populations.

Much of the mechanistic insight into protective immunity against Mtb comes from mouse models of infection. Resistant strains of mice, such as the commonly used C57BL/6J (B6), are able to restrict the replication of Mtb for over a year (13). Protective immunity in B6 mice relies heavily on Th1 biased CD4+ T cell activation and the production of IFN*γ* in the infected tissue (14, 15). IFN*γ* mediates its protective effect both by activating microbiocidal mechanisms in parasitized macrophages (16–18) and by inhibiting the recruitment of granulocytes that have been shown to exacerbate disease (19, 20). As these effects require local production of cytokine, the adhesion molecules and chemokines required for T cell recruitment play a pivotal role in immunity. Studies in knockout mice have shown that T cell expression of the integrin *α*L*β*2 and the chemokine receptors CXCR3, CCR5, and CCR2 are important for the proper positioning of these T cells and the expression of protective immunity in the lung (21–24).

Despite the wealth of mechanistic data that can be obtained in the mouse model, standard lab strains of mice do not reproduce the diversity in pathogenesis observed in natural populations. Not only does the relatively homogenous histopathology observed in these animals differ from the variable disease seen in patients (21), but recent evidence suggests an unappreciated diversity in human immune responses to Mtb (22). For example, it now appears that some humans have the capacity to control Mtb infection in the absence of the IFN*γ* response that is critical in B6 mice (23), and emerging evidence also suggests a possible protective role for antibodies that play little role in the standard mouse model (24, 25). Much of the previous work to increase the diversity of TB disease in mice has focused on relatively susceptible sub-strains that remain closely related to B6 mice (26). While studies contrasting these strains have identified a number of quantitative trait loci (QTL) associated with susceptibility (27–32), these highly-related strains still do not mimic the diversity observed in an outbred population.

Recently, a number of new resources have become available to introduce additional genetic variability into mouse model systems. For example, Diversity Outbred (DO) mice are an outbred population of genetic mosaics based on eight inbred founders, including highly divergent wild-derived strains (33). Mtb infection of DO mice produces a wide-range of disease manifestations including extreme susceptibility, which is not observed in more standard lab strains (34). While the DO population incorporates a great deal of diversity, each genotype is represented by a single unique mouse, limiting the mechanistic characterization that is otherwise a strength of the mouse system. The Collaborative Cross (CC) consists of recombinant inbred lines derived from the same eight inbred founder strains as the DO (35). In contrast to the outbred DO, each CC strain represents a reproducible mosaic of the founder genomes, and thus a reproducible model of disease (36). We have shown that the range TB susceptibility observed in the DO population can be recapitulated in CC mice (37). This previous work found that CC042/GeniUnc (CC042) animals was highly susceptible to Mtb infection, in contrast to resistant strains such as CC001/Unc or B6 (37).

Here we investigated the basis TB susceptibility in the CC042 strain using a parallel genetic and immunophenotyping approach. By producing an intercross population based on CC042 and the resistant CC001 strains, we identified multiple QTL, named “Tuberculosis ImmunoPhenotype” (*Tip1-4*), that were differentially associated with bacterial burden and/or IFN*γ* production. In this population, canonical IFN*γ*-dependent immunity was controlled by a novel mutation in the *Itgal* gene, which disrupts expression of the *α*L*β*2 adhesion molecule and prevents the recruitment of cytokine expressing T cells to the site of infection. Other *Tip* loci were driven by wild-derived founder alleles that either reduce IFN*γ* production or control IFN*γ*-independent immunity. Together, these observations explain the extreme susceptibility of CC042 and indicate that the CC panel can be used to understand diverse mechanisms of protective immunity to Mtb.

## Results

### Control of Mtb infection in CC042 is lost upon the onset of adaptive immunity

In order to dissect the mechanisms underlying CC042 susceptibility to Mtb, we first profiled the kinetics of disease in aerosol-infected CC042 animals compared to more resistant B6 mice. In the standard B6 model, the peak of bacterial burden was observed around 21 days post infection, coincident with the onset of robust Th1 immunity. While CC042 had 10-fold lower CFU in the lung and spleen at 14 days, compared to B6 (Fig1A, B), CC042 animals ultimately failed to control bacterial replication. By day 28 post-infection, the lungs of CC042 mice harbored 100-fold more bacteria compared to B6. Similar trends in relative bacterial burden were observed in the spleen. All CC042 mice had lost significant weight and required euthanization because of morbidity by day 33 post infection (Fig1C, D). Both male and female CC042 mice were similarly moribund at this timepoint.

The superior control of bacterial replication by B6 mice correlated with IFN*γ* abundance in lung homogenate. There was a significant increase in IFN*γ* levels starting on day 21, which peaked by day 28 in all B6 mice (Fig1E). In contrast, the concentration of IFN*γ* in the lungs and spleens of CC042 mice remained relatively low throughout the course of infection (Fig1E, F). Further histological comparison found a cellular infiltration in B6 lung lesions that largely consisted of macrophages and lymphocytes throughout the experiment, while necrosis and neutrophil infiltration was apparent in CC042 lungs by 21 days, and was sustained until termination of the experiment (28 – 33 days post infection) (Fig 2). Together, these data suggested that the susceptibility of CC042 could be related to a defect in IFN*γ* production that promoted bacterial growth and granlulocyte infiltration.

### Identifying TB susceptibility loci in a [CC001xCC042] intercross

To investigate the genetic basis of CC042 susceptibility, we took an F_2_ intercross approach to segregate TB disease traits and map the causal genetic variants. CC001 was chosen as the CC cross partner because of its relative TB resistance (similar to a B6) (37) and to match the CC042 H-2^b^ locus. We crossed female CC001 with male CC042 mice to generate F_1_ progeny (CC001xCC042)F_1_, which were then intercrossed to produce ∼200 F_2_ offspring. We infected male and female F_1_ and F_2_ progeny, along with parental strains, with Mtb (H37Rv) via low-dose aerosol. Mice were sacrificed between 28-31 days post-infection, a timepoint that maximized phenotypic differences, while minimizing morbidity. Measured phenotypes included lung colony forming units (CFU), spleen CFU, and lung IFN*γ* levels.

For spleen CFU and lung IFN*γ* levels, F_1_ mice showed an intermediate phenotype, and F_2_ mice displayed a distribution of values spanning the range of the parental strains (Fig 3A, B). In contrast, overdominance was observed for lung CFU, as F_1_ mice had higher bacterial burdens in the lung than the susceptible CC042 mice (Fig 3C) and F_2_ mice spanned this greater phenotypic range. The three traits co-varied in a predictable manner (Fig 3D). Bacterial burden in lung and spleen were positively correlated. Lung IFN*γ* levels were more strongly associated with CFU in spleen than lung, consistent with the more prominent role of IFN*γ*-independent T cell functions in the lung (38). The imperfect correlation between these traits, as well as the expansion of phenotypic ranges in F_2_ animals, suggested that multiple genes controlled the differences between CC001 and CC042.

In total, 169 F_2_ mice were genotyped with the MiniMUGA array (39). We first validated our genetic mapping protocol using a coat color trait (Supp Fig 1). CC042 have a white head spot (“blaze”) on their forehead, a characteristic inherited from the WSB/EiJ (designated for Watkins Star Blaze; WSB) founder strain. One in every four F_2_ progeny carried a blaze, confirming autosomal recessive inheritance (40). The blaze and base coat color (black from CC001 or agouti from CC042) assorted independently in the F_2_ offspring, as shown by the expected 9:3:3:1 ratio (Supp Fig 1). QTL mapping on the presence or absence of white head spotting in the F_2_ mice identified a highly significant QTL on Chromosome 10 (Supp Fig 1). This interval contained kit ligand (*Kitl*, stem cell factor), which was previously shown to underly this trait in the pre-CC population (40).

**Figure 1.**
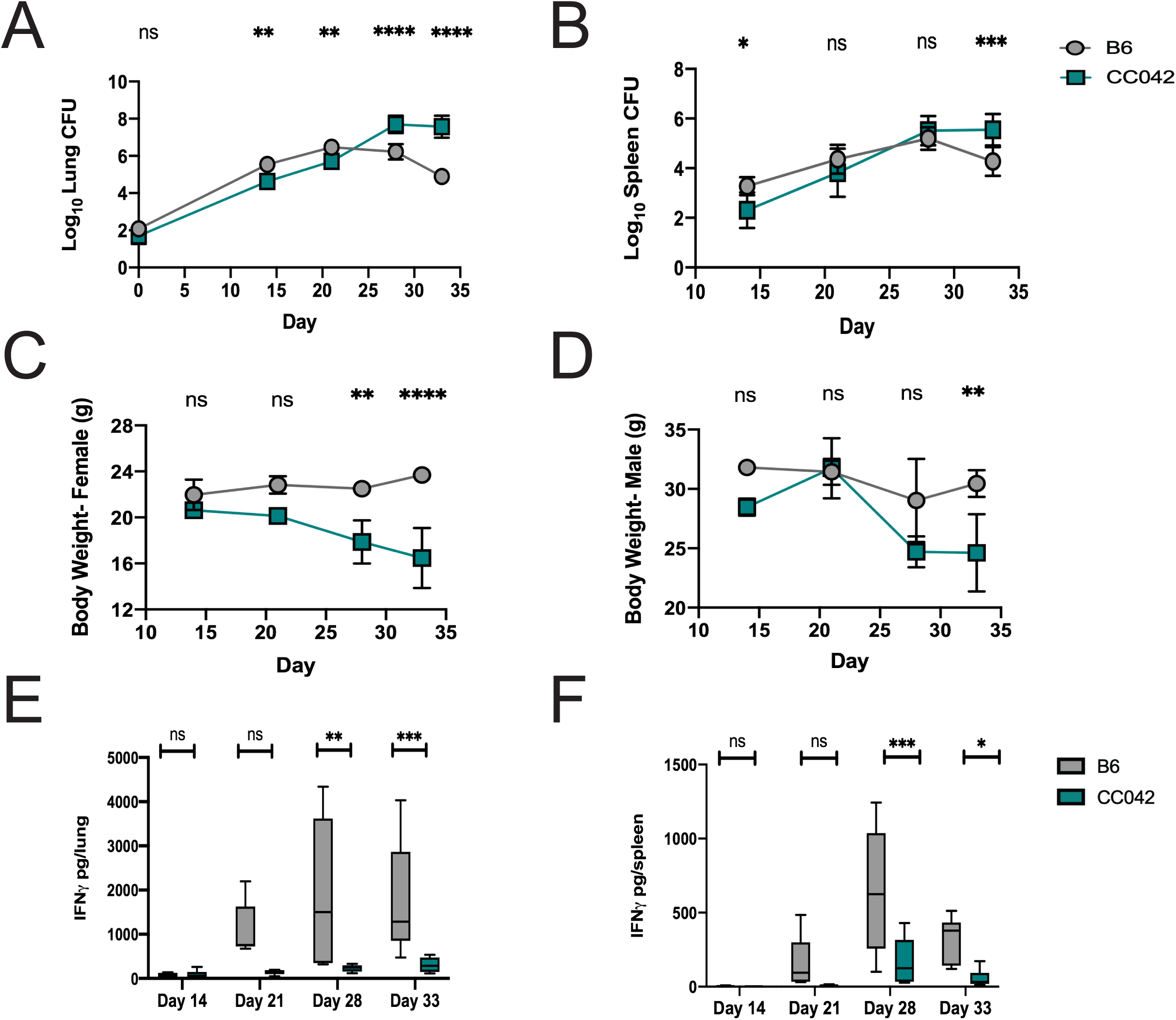
CC042 are susceptible to low-dose aerosol Mtb infection. Lung CFU (A), Spleen CFU (B), body weight (C, D), and total IFN*γ* levels in lung (E) or spleen (F) homogenate at 14, 21, 28, and 33 days post infection by low-dose aerosol (∼50-100 CFU) of *Mtb* strain H37Rv. All mice were infected in one batch and 3 males and 3 females for each strain were used for analysis at each time point. Graphs represent mean ± SD. One-way ANOVA with Sidak’s multiple comparisons test was used to determine significance, where p<0.05 *, p<0.01**, p<0.001***, and p<0.0001****.

Next, we conducted QTL mapping on the tuberculosis-associated phenotypes Lung CFU, Spleen CFU and IFN*γ* levels. Using batch and sex as covariates, we identified four significant QTL that affect measured Tuberculosis ImmunoPhenotypes (*Tip1-4*) (Table 1 and Figure 4A, B).

**Figure 2.**
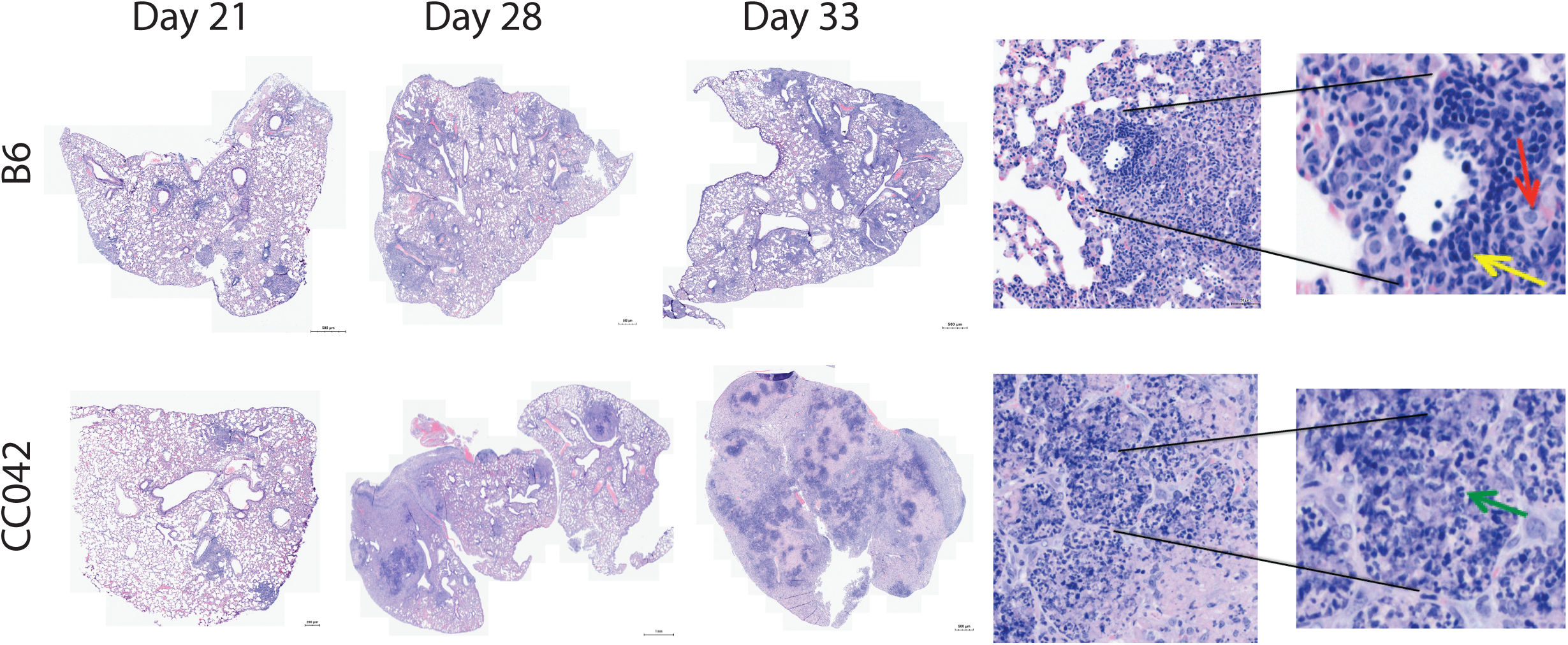
Changes in lung pathology during Mtb infection. Hematoxylin and eosin staining of lung lobes from B6 and CC042 mice at Day 21, 28, and 33 post infection. Images are representative of 6 mice per strain per time point. Magnified insets from D33 images show lymphocytes (yellow arrow), macrophages (red arrow) and neutrophils (green arrow).

**Figure 3.**
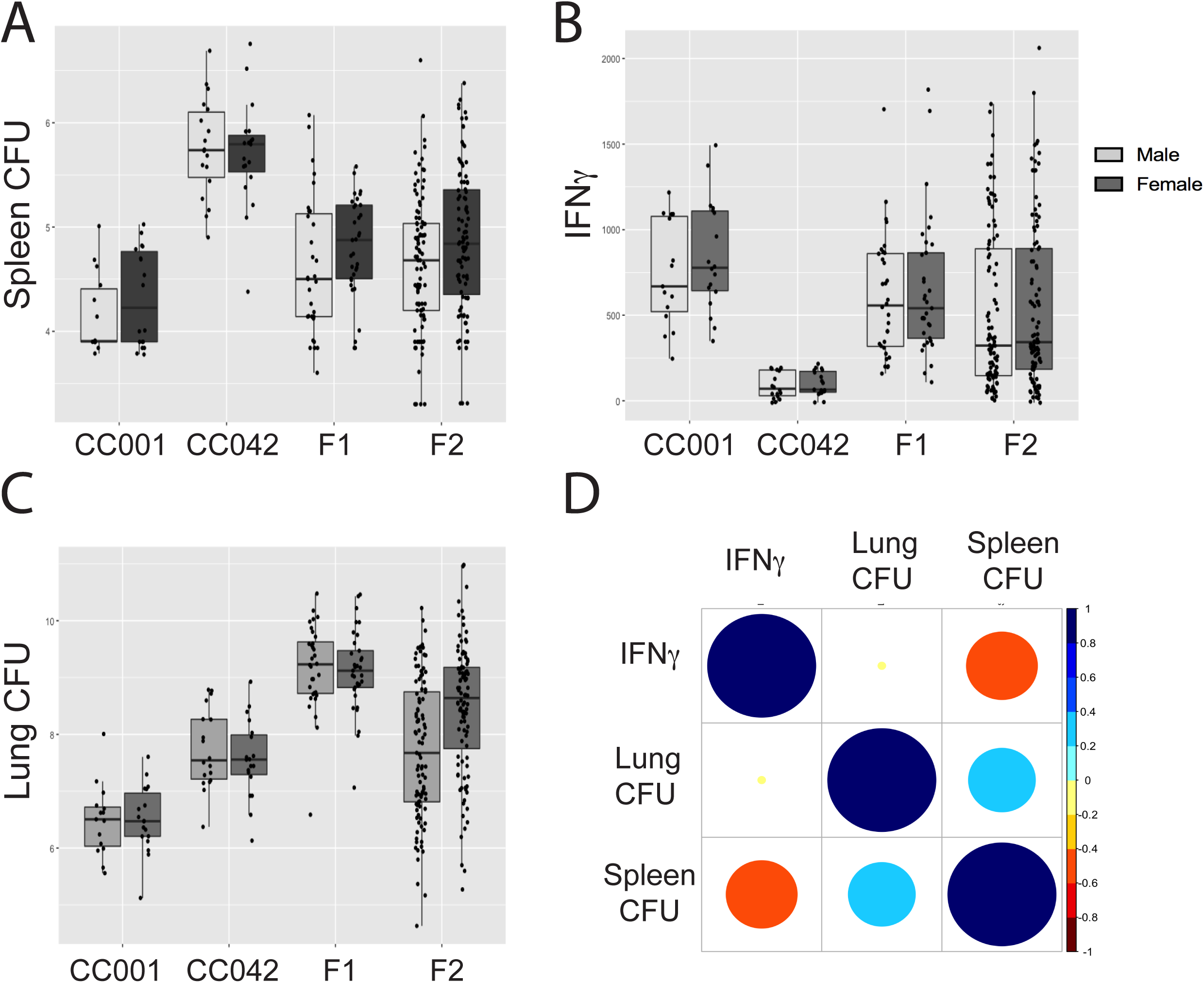
TB disease traits in a CC001 × CC042 F_2_ intercross population. At 28-31 days post infection the following traits were quantified in parental strains, and F_1_ and F_2_ offspring: (A) Lung CFU (B), spleen CFU (C) and total IFN*γ* from lung homogenate. The Pearson correlation between measured traits is shown in (D). Data shown are from the following population sizes: F_2_ (n = 201), F_1_ (n = 65), CC001 parent (n = 33) and CC042 parent (n = 37). Mice were infected in 4 batches and values were adjusted for batch differences using coefficients from multiple regressions.

**Figure 4.**
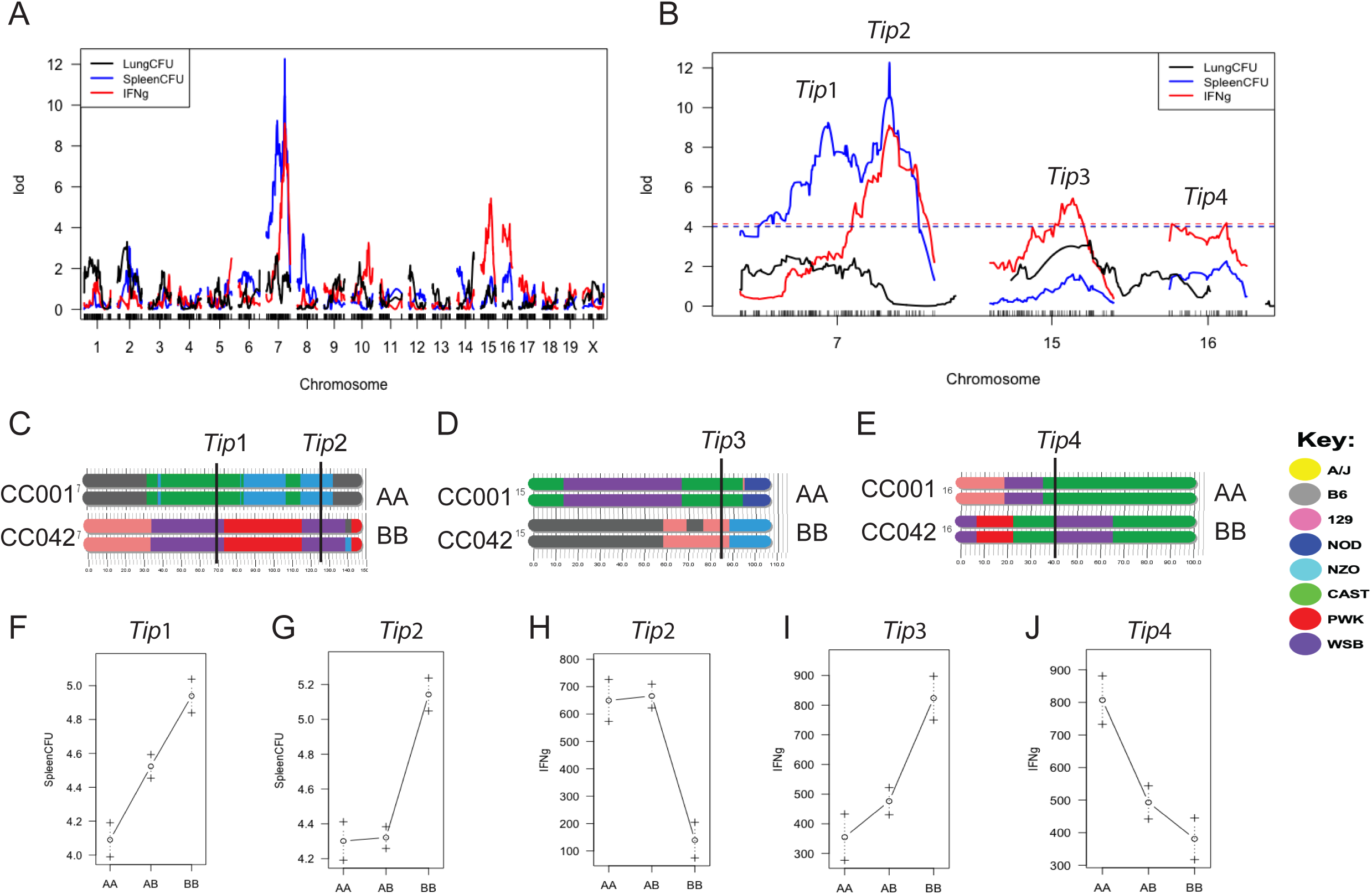
QTL mapping identifies four loci underlying TB susceptibility. QTL scan of lung CFU, spleen CFU, and IFN*γ* abundance in the lung identify four Tuberculosis immune phenotype (*Tip*) loci on Chromosomes 7, 15, and 16 (A, B). Dotted lines in B represent 5% false discovery rate for each trait based on permutation analysis. *Tip* loci on each chromosome are indicated at the marker with peak LOD. (F-J) Allele effect plots for the indicated *Tip* loci. One hundred seventy (170) F_2_ mice were successfully genotyped and used for QTL mapping.

**Table 1.** Tuberculosis ImmunoPhenotype (*Tip*) QTL in an CC001xCC042 F2 cross. Low and High haplotypes are provided at the peak logarithm of the odds (LOD) of each QTL. P value was determined by permutation test. High and low haplotypes are provided relative to each specific trait. Fraction of variance explained by each QTL (Variance (%)) was estimated by fitting a single QTL model for each trait at the respective peak locations with sex and batch as covariates.

| QTL | Chr | Trait | P value | Max LOD | Peak marker | Peak (Mb) | Bayes Interval (Mb) | Low haplotype | High haplotype | Inheritance mode | Variance (%) |
| --- | --- | --- | --- | --- | --- | --- | --- | --- | --- | --- | --- |
| Tip1 | 7 | Spleen CFU | < 10<sup>-4</sup> | 9.2 | gUNC13104259 | 72.1 | 3.6 - 72.9 | CAST | WSB | Additive | 17.6 |
| Tip2 | 7 | Spleen CFU | < 10<sup>-4</sup> | 12.3 | gUNC13793270 | 125.4 | 124.7 -127.3 | NZO | WSB | recessive | 22.5 |
| Tip2 | 7 | IFNγ level | 7 x 10<sup>-4</sup> | 9.1 | gUNC13793270 | 125.4 | 121.0 - 130.7 | NZO | WSB | recessive | 21.4 |
| Tip3 | 15 | IFNγ level | 0.011 | 5.4 | mbackupUNC150396514 | 83.1 | 53.7 - 89.7 | CAST | 129 | Ambiguous | 13.5 |
| Tip4 | 16 | IFNγ level | 0.046 | 4.2 | UNC26693650 | 40.8 | 4.5 - 44.7 | WSB | CAST | Ambiguous | 10.6 |

Spleen CFU mapped to two distinct QTL on Chromosome 7; proximal *Tip1* at 72Mb and distal *Tip2* at 125Mb (Figure 4C). *Tip1* was inherited in an additive fashion, with heterozygous mice carrying both CAST/EiJ (CAST) and WSB haplotypes exhibiting an intermediate phenotype relative to both homozygotes (Fig 4F). In contrast, the susceptibility phenotype associated with *Tip2* was recessive, with mice homozygous for the WSB allele demonstrating a tenfold increase in spleen CFU on average (Fig 4G). IFN*γ* production in the lung was associated with 3 distinct QTL. The main locus explained 21.4% of the variation and was mapped to the *Tip2* region on chromosome 7 (Fig 4B), evidence that the same variant likely controls IFN*γ* and bacterial burden at this locus. Two additional QTL were also associated with IFN*γ* levels, mapping to Chromosomes 15 and 16 (*Tip3* and *Tip4* respectively) (Fig 4A and B). A second potential QTL on proximal Chromosome 16 was also associated with IFN*γ*, however its LOD did not reach genome wide significance at P<0.05. At *Tip3*, low IFN*γ* levels were associated with haplotypes from the CAST (Fig 4D) founder, a strain previously found to lack IFN*γ* expression in the lung upon infection with either Mtb or pox virus (37, 41, 42). Notably, *Tip1* was associated with CFU, but not IFN*γ*, indicating that this variant was functionally distinct from *Tip2*. No QTL were associated with lung CFU, suggesting that this trait is under more complex genetic control than the others. In sum, these functionally and genetically diverse QTL indicated that the immune response to Mtb was under multigenic control in these strains.

Considering that *Tip1* and *Tip2* are both on Chromosome 7 and are driven by the WSB parent haplotype, we tested the independence of these peaks by remapping spleen CFU using the genotypes at *Tip1*as a covariate. After removing the variation explained by the proximal *Tip1* QTL, the distal *Tip2* QTL still met the threshold for genome-wide significance (Supp Fig 2A). In addition, fitting a three QTL model to the spleen CFU phenotypes showed that *Tip1*, *Tip2*, and *Tip3* all contributed additively; removing any of them from the full model resulted in a significantly poorer fit (Supp Fig 2B). Altogether, we identified four independent QTL in the F_2_ cross, indicating multigenic control of Mtb immunity in this F_2_ population.

### T cell function and recruitment are impaired in CC042 mice

Concurrent with our genetic mapping strategy, we enumerated changes in leukocyte cell types that could alter the susceptibility of CC042 relative to B6 mice. Accumulation of T and B lymphocytes in the lungs of B6 mice began between 14 and 21 days post-infection (Fig 5A, B). In contrast, the numbers of T and B cells in the lungs of CC042 mice were significantly reduced. This paucity of lymphocyte accumulation in the lungs of CC042 mice was mirrored a dramatic increase in CD11b+Gr1+ granulocytes (Fig 5C), consistent with the neutrophilic infiltrates observed in the lung histopathology (Figure 2). Significant differences in the CD11b+Gr1-monocytes/macrophage subset between these mice was only apparent at the last timepoint, at which time the CC042 animals had become moribund (Fig 5D).

**Figure 5.**
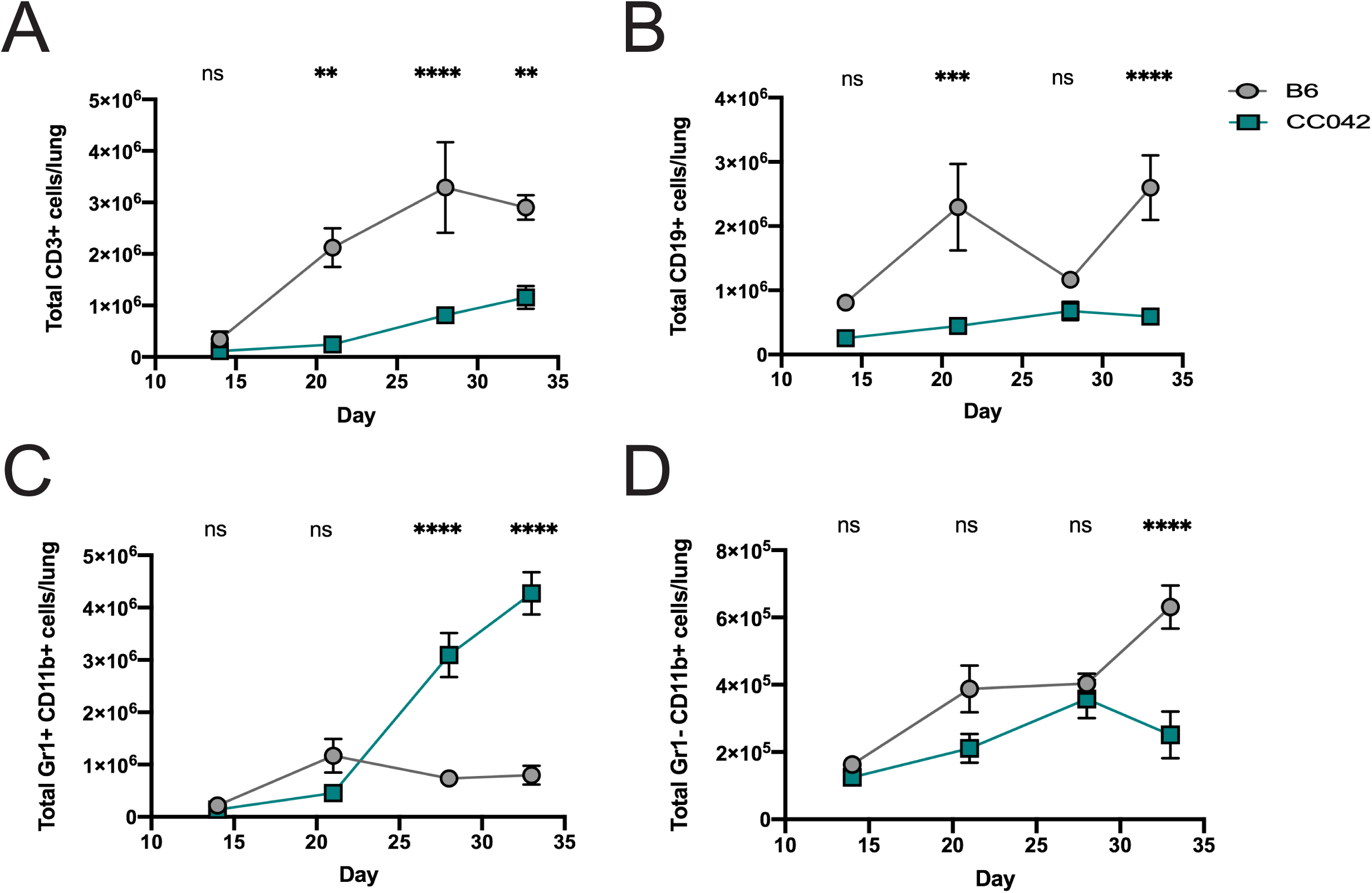
Susceptibility of CC042 correlates with altered numbers of lung leukocytes. Total number of the following cells were enumerated in the lungs of B6 and CC042 mice: (A) T cells (lymphocytes > single cells > CD3+ CD19-), (B) B cells (lymphocytes > single cells > CD3-CD19+), (C) neutrophils (lymphocytes > single cells > Gr1+ CD11b+), and (D) monocytes/macrophages (lymphocytes > single cells > Gr1-CD11b+). All mice were infected in one batch and 3 males and 3 females for each strain used for analysis at each time point. Graphs represent mean ± SD. One-way ANOVA with Sidak’s multiple comparisons test was used to determine significance where p<0.05 *, p<0.01**, p<0.001***, and p<0.0001****.

The reduction in IFN*γ* production and paucity of pulmonary T cells in the lungs of CC042 mice suggested that their susceptibility to Mtb might be related to a defect in T cell function. To assess effector function, we isolated cells from the lungs of infected B6 and CC042 mice and stimulated them with anti-CD3 to determine whether T cells from CC042 mice had differentiated into a distinct T cell subset (e.g. Th1 vs. Th2, Th17, Treg). Using a standard intracellular cytokine staining (ICS) approach, we found that CD4 and CD8 T cells from CC042 mice produced less IFN*γ*, TNF, IL-17a, IL-2, IL-10 and CD107a upon stimulation in comparison to B6 T cells (Fig 6A, B). Thus, instead of representing an altered T cell differentiation state, the lack of cytokine production by T cells from the lungs of infected CC042 mice may indicated that there is an impairment in either T cell priming or recruitment of T cells to the lung.

**Figure 6.**
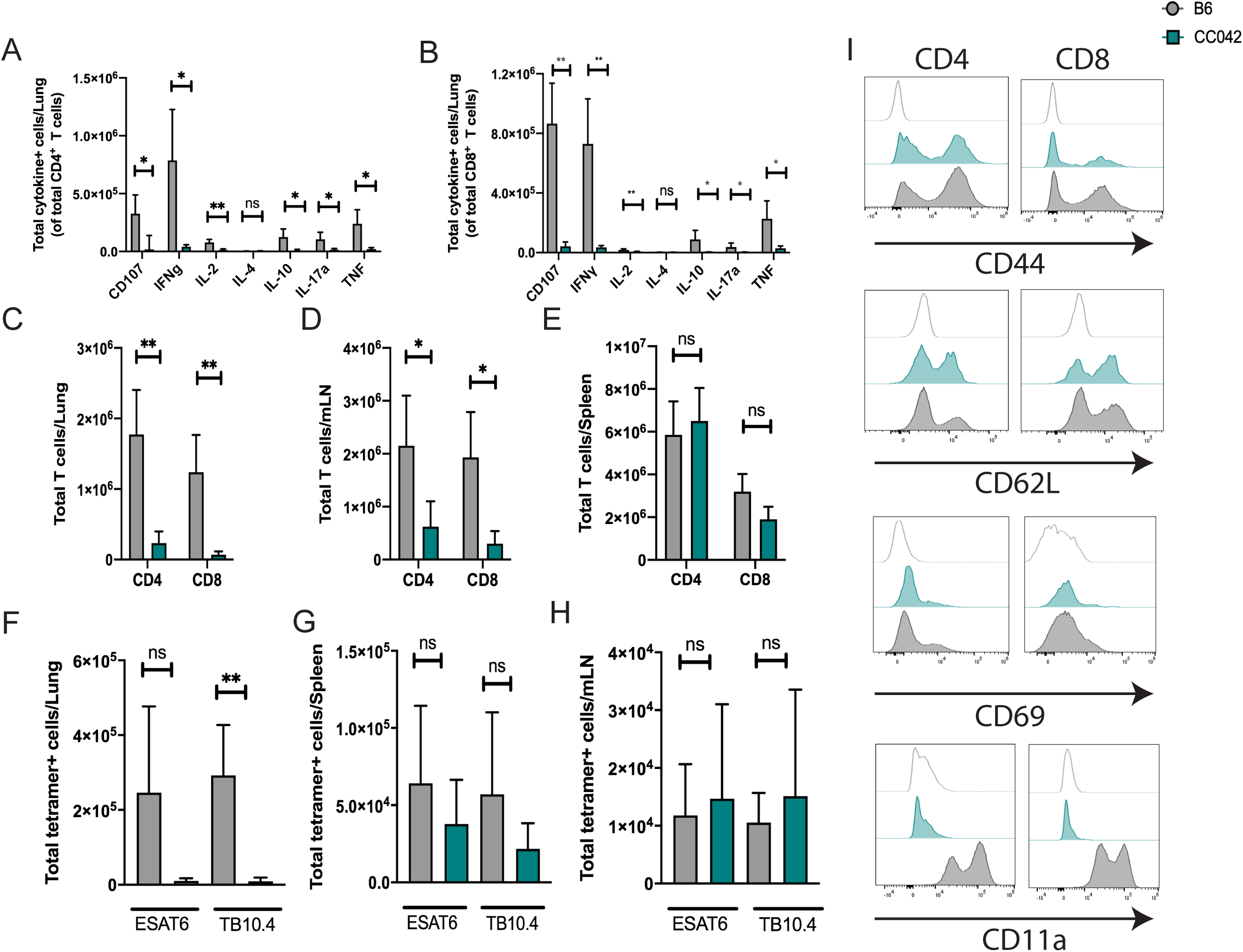
CC042 have a defect in T cell recruitment to the lung and lack CD11a expression. Intracellular cytokine staining (ICS) of CD4 (A) and CD8 (B) T cells reveals a defect the number of cytokine-producing T cells in the lungs of CC042 mice. Total number of CD4 and CD8 T cells in the lung (C), mediastinal lymph node (D), and spleen (E). Total number of ESAT6 (CD4) or TB10.4 (CD8) tetramer positive cells in the lung (F), mediastinal lymph node (G), and spleen (H). Bar plots show mean + SD. Welch’s t-test was used to determine significance where p<0.05 *, p<0.01**. Histograms (I) of CD4 (left) and CD8 (right) T cells stained for activation and migration markers CD44, CD62L, CD69, and CD11a. Isotype control (top, light gray trace), CC042 (middle, teal), and B6 (bottom, gray).

In order to distinguish between these possibilities, we counted the number of CD4 and CD8 T cells in the lung, mediastinal lymph node (mLN) and spleen at 28 days post infection using flow cytometry. At the same time, we enumerated the number of antigen specific CD4 and CD8 T cells using ESAT-6 (CD4) and TB10.4 (CD8) tetramers. CC042 mice had fewer total CD4 and CD8 T cells in the lung compared to B6 mice (Fig 6C). This was also true in the mLN but not the spleen (Fig 6D, E). While we saw that CC042 mice had significantly fewer ESAT-6-specific CD4 and TB10.4-specific CD8 T cells in the lung than B6 animals, both groups of mice had similar numbers of antigen-specific T cells in the mLN and the spleen (Fig 6 F, G, H). When antigen-specific T cells were considered as a fraction of total T cells, the frequencies of ESAT-6-specific T cells was similar in the lung, spleen and mLN of B6 and CC042 mice, while the frequency of TB10.4-specific T cells were lower in the lungs of CC042 mice but were higher in the spleen and mLN in comparison to B6 mice (Supp Fig 3). Taken together, this suggests that T cell priming of Mtb-specific CD4 and CD8 T cells was occurring in the draining LN, and the diminished T cell numbers in the lungs of CC042 mice is due to an impairment in T cell recruitment. As such, we examined the cell surface expression of CD44, CD69, CD62L and CD11a on CD4 and CD8 T cells from the lungs of infected CC042 and B6 mice, as these markers have been classically associated with either T cell activation or migration (41, 42). We found that CD4 and CD8 T cells from the lungs of CC042 and B6 mice had appropriately upregulated CD44 and CD69 and downregulated CD62L (Fig 6I). However, CD11a was undetectable on both CD4 and CD8 T cells from CC042 mice. As CD11a is the *α*L component of *α*L*β*2, the principal β2-integrin on T cells that is crucial for lymphocyte trafficking, this defect could explain many of the immunological differences observed between CC042 and B6.

### A CC042 private mutation in Itgal explains Tip-2-driven susceptibility

The gene encoding CD11a, *Itgal*, is located on chromosome 7, within the *Tip2* locus identified in our intercross. The lack of CD11a expression on CC042 lymphocytes implicated *Itgal* variation as the basis for *Tip2*. To investigate this possibility, CD11a expression was assayed in WSB and CC011 strains, which contain the susceptibility-associated WSB haplotype at *Tip2* (Fig 7a). We found that WSB and CC011 splenocytes expressed similar CD11a levels as B6 mice, leading us to hypothesize that CC042 had incurred a private mutation during inbreeding that impacts CD11a production.

**Figure 7.**
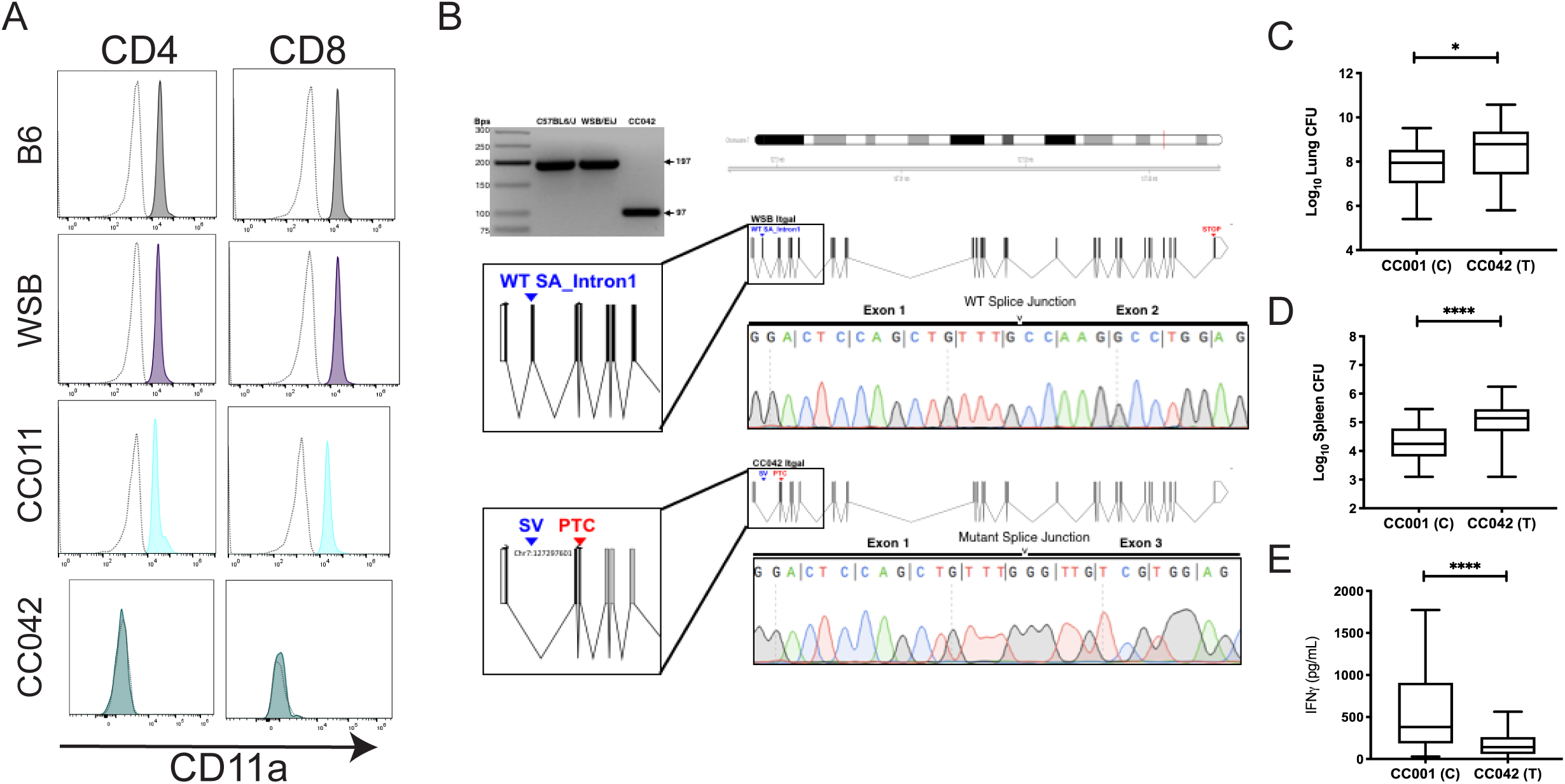
A CC042 private mutation in *Itgal* explains *Tip2* driven susceptibility. (A) Histograms depicting CD11a staining in CD4 (left) and CD8 (right) T cells from B6 (gray shading), WSB (purple shading), CC011 (light blue shading), and CC042 (teal shading). The isotype control antibody staining is shown on each plot as a dotted gray trace. (B). PCR primers flanking the putative private *Itgal* mutation were used to amplify cDNA from B6, WSB, and CC042 RNA. PCR products were separated by gel electrophoresis on a 3% agarose gel. Lanes 2 and 3 show the 197bp amplicons for B6 and WSB, the parental allele for CC042. The 100bp size decrease in the CC042-derived product is consistent the loss of exon 2 (lane 4 and schematic). Sanger sequencing traces from the CC042 and WSB amplicons are shown. TB immunophenotypes were reevaluated in F_2_ progeny of the CC001xCC042 that were homozygous for each parental allele at the *Itgal* locus (probe UNC13811649): (C) Lung CFU, spleen (D) CFU, and (E) lung IFN*γ* levels Welch’s t-test was used to determine significance where p<0.05 *, p<0.01**, p<0.001***, and p<0.0001****. Box and whiskers plots indicate the median and min to max values.

Using whole genome sequences of representative CC strains, variants private to each have been identified (43, 47). Using this dataset, we found that the CC042 genome sequence contains a 15 bp deletion in the first intron of the Itgal1 gene, which is not present in the ancestral WSB allele. This deletion alters the canonical splice acceptor sequence (“AG”) at the 3’ terminus of intron 1 (http://Jul2019.archive.ensembl.org/Mus_musculus_WSB_EiJ/Transcript/Exons?db=core;g=MGP_WSBEiJ_G0032170;r=7:131372531-131430694;t=MGP_WSBEiJ_T0085398). Although the resultant mutant sequence (“TG”) has been observed to function as a splice acceptor in certain transcripts (43), we hypothesized that this mutation could alter splicing. Amplification and sequencing of a fragment spanning exons 1 and 3 of the *Itgal* mRNA confirmed that the CC042 transcript lacked the exon 2 sequence that was contained in both the B6 and WSB mRNAs (Fig. 7B). The lack of exon 2 is predicted to produce a frameshift and premature termination (Fig. 7B). While *Tip2* was only significantly associated with spleen CFU and lung IFN*γ* in the whole genome scans, the lack of T cells in the lungs of CC042 animals suggested the CD11a deficiency also influenced bacterial replication at this site. To test this hypothesis, we used the measured phenotypes from the F_2_ cross and found that the CC042 *Itgal* allele was significantly associated with higher CFU values (Fig7 C-E). This analysis indicated that the *Itgal* mutation affects all metrics of tuberculosis susceptibility in CC042.

## Discussion

In this work, we used a classic genetic strategy to investigate the diversity of responses to Mtb infection observed in CC strains. Previous work identified a number of genetically-dissociable immunophenotypes in Mtb-infected CC founder lines, including bacterial load and IFN*γ* production (37). The intercross between two phenotypically divergent CC strains reported here supports the independence of these traits and demonstrates that multiple phenotypes can be mapped simultaneously using this strategy. Investigating the genetic architecture of TB disease in these highly diverse mice revealed a number of new insights that would not have been apparent in intercrosses between more genetically-homogenous lab strains.

A loss-of-function mutation in *Itgal* was found to account for the *Tip2* QTL and explain a significant portion of the susceptibility of the CC042 line. We previously found that *Itgal* deletion in the B6 background resulted in a defect in T cell recruitment to the lung and increased pulmonary Mtb burden (44), both traits that are associated with *Itgal*-deficiency in the CC001xCC042 intercross. While *Tip2* was only significantly associated with spleen CFU and IFN*γ* production in whole genome scans, *Itgal* genotype also correlated with lung CFU when examined in isolation. Thus, we conclude that *Itgal* affects pathogenesis in both lung and spleen, and the lack of genome wide association between *Itgal* and lung CFU is likely due to complex genetic factors, such as the observed overdominance and the possible presence of additional variants that dilute the effect of the *Itgal* mutation.

While the disease-promoting effect of CD11a deficiency was generally consistent between the B6 background and the recombinant CC genotype, the loss of *Itgal* is insufficient to explain the susceptibility of the CC042 strain. These CC042 mice succumb to infection approximately five months earlier than *Itgal^−/−^* B6 mice (44), likely due to the more dramatic infiltration of granulocytes and necrosis observed in the lung. The increased susceptibility of CC042 animals can be attributed, at least in part, to *Tip*1. The variant(s) underlying *Tip1* appear to be functionally distinct from *Itgal*, as the effect of *Tip*1 and *Tip*2 are additive, these traits differ in their mode of inheritance, and *Tip1* is not associated with IFN*γ* levels. These data indicate that IFN*γ*-independent mechanisms, such as those underlying *Tip*1, can act in an additive fashion to increase the susceptibility of animals with a more canonical immunodeficiency that affects Th1 cell activity.

This intercross between two very diverse genotypes allowed the mapping of QTL that may be associated with an additional trait previously identified in the CAST founder strain. Despite being relatively resistant to Mtb infection, CAST mice do not produce detectable levels of IFN*γ* in the lung (37). This observation is one of many indicating that IFN*γ*-independent immune mechanisms play an important protective role, particularly in the lungs (38, 45, 46). *Tip1*, and *Tip*3, are likely related to the IFN*γ*-deficient phenotype of CAST mice. This founder haplotype at *Tip3* is associated with low IFN*γ* production, and the CAST allele at *Tip1* reduces CFU without influencing IFN*γ* levels. Defining the basis of this phenotype, might represent an important step to understanding the immune response to Mtb in the recently identified subset of humans that control Mtb infection in the absence of a detectable IFN*γ* response (23).

The CC population was initially envisioned as a genetic mapping resource, based on the random distribution of founder alleles between strains (47). While the CC panel been shown to be valuable for this type of study, the *Itgal* mutation identified in this work was not derived from a founder line and likely occurred during the process of inbreeding CC042. Such mutations in individual CC strains provide an additional advantage to studies within the CC: both common variants, as well as private variants (∼28,000) circulate in this population and can drive disease responses (39, 48). While these mutations are invisible in genetic association studies that are based on comparison of CC lines, our work shows that their effect can be revealed through intercrosses and the molecular characterization of these relatively rare variants can be rapidly achieved. This feature, in combination with the variety of phenotypes that can be addressed in CCxCC intercrosses, highlight the value of this approach.

The immune response to Mtb in natural populations is variable. Several lines of recent evidence suggest the importance of mechanisms distinct from canonical Th1 immunity that dominate in the classic “mouse model” of TB, which relies on a small number of genetically similar mouse lines. Using a simple intercross strategy, we leveraged the genetic diversity of CC lines to define chromosomal loci controlling three distinct TB-related traits; *Itgal*-dependent T cell recruitment (*Tip*2), IFN*γ*-independent bacterial control (*Tip*1), and IFN*γ* production (*Tip*3-4). All of these traits associated with haplotypes that are absent from standard mouse strains, supporting the value of genetic diversity to understand highly variable traits, such as TB susceptibility.

## Materials and Methods

### Ethics statements and experimental animals

C57BL/6J (#0664) (B6) mice were purchased from the Jackson labs. CC042/GeniUnc and CC001/Unc mice were obtained from the Systems Genetics Core Facility at the University of North Carolina (49), and bred at UMASS Medical School under specific pathogen-free conditions and in accordance with the University of Massachusetts Medical School IACUC guidelines. F_1_ mice were generated from crossing CC001 females by CC042 males (CC001xCC042). The F_2_ mice used for QTL mapping were obtained from crossing these F_1_ mice (e.g. all F_2_ animals were ((CC001xCC042)x(CC001xCC042))-F_2_.

### Mycobacterium tuberculosis infection

Wild type *M. tuberculosis* strain H37Rv (PDIM positive) was used for all studies unless indicated. Prior to infection, bacteria were cultured in 7H9 medium containing 10% oleic albumin dextrose catalase growth supplement (OADC) enrichment (Becton Dickinson) and 0.05% Tween-80. For aerosol infections, bacteria were resuspended in phosphate-buffered saline containing Tween-80 (PBS-T). Prior to infection, bacteria were sonicated then delivered via the respiratory route using an aerosol generation device (Glas-Col). Groups of mice were sacrificed 24 hours post-infection to enumerate infectious dose. The infectious dose for all experiments ranged from 50-150 CFU. Both male and female mice were used throughout the study, as indicated. All animals used for experiments were 8-12 weeks old.

### CFU enumeration and cytokine quantification

To determine CFU, mice were anesthetized via inhalation with isoflurane (Piramal) and euthanized via cervical dislocation. The organs were aseptically removed and individually homogenized, and viable bacteria enumerated by plating 10-fold serial dilutions onto 7H10 agar plates. Plates were incubated at 37°C, and colonies were counted after 21 days. Cytokine concentrations in cell-free lung homogenates were quantified using commercial enzyme-linked immunosorbent assay (ELISA) kits (IFN*γ* Duo Set #DY485 R&D Systems) according to manufacturer instructions.

### Histology

Lung lobes from B6 and CC042 mice infected with *Mtb* were fixed in 10% neutral buffered formalin, embedded in paraffin, and sectioned at 5μm. Sections were stained with hematoxylin and eosin (H&E). All sectioning and staining was done by the Diabetes and Endocrinology Research Center Morphology Core (DERC) at the University of Massachusetts Medical School. Images were captured on a TissueGnostics TissueFAXS PLUS at 2X and 20X magnification.

### Flow cytometry analysis

Lung tissue was harvested in RPMI containing FBS and placed in C-tubes (Miltenyi). Collagenase type IV/DNase I was added and tissues were dissociated for 10 seconds on a GentleMACS system (Miltenyi). Tissues were incubated for 30 minutes at 37°C with oscillations and then dissociated for an additional 30 seconds on a GentleMACS. Lung homogenates were passed through a 70-micron filter. Cell suspensions were washed in RPMI, passed through a 40-micron filter and aliquoted into 96 well plates for flow cytometry staining. Non-specific antibody binding was first blocked using Fc-Block, after which cells were then stained with CD3-BV785 (145-2CL1), CD8-APC-Fire 750 (53-8.7), CD44-PerCP-Cy5.5 (IM7), CD11a-BV711 (M17/4), CD69-PE-Cy7 (H1.2F3) and CD62L-BV570 (MEL-14) from Biolegend and CD4-Alexa Fluor 700 (RM4-5) from BD Biosciences. In some experiments, intracellular cytokine staining was also performed. After surface marker staining, cells were subsequently permeabilized using cytofix/cytoperm (BD Biosciences) and stained with IFN-APC (XMG1.2), IL-2 PE-Cy7(JES6-5H4), IL-17a-BV650 (TC11-18H10.1), TNF-BV421 (MP6-XT22), IL-10 (JES5-16E3) and CD107a-PE (1D4B) from Biolegend and IL-4-Alexa Fluor 488 (11B11) from Invitrogen. Live cells were identified using fixable live dead aqua (Life Technologies). Cells were stained for 30 minutes at room temperature and fixed in 1% Paraformaldehyde for 60 minutes. All flow cytometry was run on either a MACSQuant Analyzer 10 (Miltenyi) or Aurora (Cytek) and was analyzed using FlowJo_V10 (Tree Star).

### Genotyping and QTL mapping

1500 nanograms of DNA was genotyped by Neogen Inc. using the the MiniMuga array. We filtered markers to those that were consistent within a previously published set of CC042 and CC001 genotypes (39), and diagnostic between these strains (*i.e.*, CC001 genotype ≠ CC042 genotype and F_1_ genotype called heterozygous). After finding and removing misplaced markers, identified using the droponemarker function of the R package qtl (R/qtl), regions of dense marker coverage were thinned to a spacing of 0.1 cM. The final genetic map contained 1806 markers.

Genotype and phenotype data were imported into R (version 3.4.3) and reformatted for R/qtl (version 1.42-8). Genotype probabilities were calculated at 0.25 cM spacing, and QTL mapping was carried out using the scanone function using batch and sex as additive covariates. Significant LOD thresholds were established by permutation test with 10,000 permutation replicates. Multi-QTL models were fit using R/qtl’s fitqtl function. LOD profiles and effect plots were generated using the plotting functions of the R/qtl package.

### Ex vivo RNA isolation and qRT-PCR

Bone marrow derived macrophages (BMDMs) from C57BL/6J, WSB/EiJ and CC042 mice were generated. Briefly, marrow from femurs and tibia of age- and sex-matched mice was isolated and cultured in high glucose DMEM (Gibco 11965092) supplemented with L-glutamine, 10% fetal bovine serum (FBS, Sigma F4135), and 20% L929 conditioned media. After seven days, differentiated cells were lifted with PBS with 10mM EDTA and seeded for subsequent experimentation. RNA was isolated by lysing the cells in Trizol (ThermoFisher 15596018) and purified using Direct-zol RNA Miniprep plus kit (Zymo Research, R2070) per the manufacturer’s recommendations. Following RNA quantification by Nanodrop, samples were diluted to 5ng/uL and used for qRT-PCR with Luna One-Step Universal qPCR kit (NEB E3005). Gene specific primers for target transcripts were used at a final concentration of 400uM with 15ng of RNA. Primer sequences for B-actin (RT-Actb_1F 5’ – GGCTGTATTCCCCTCCATCG – 3’ and RT-Actb_1R 5’–CCAGTTGGTAACAATGCCATGT–3’) and Itgal (RT-Itgal_1F 5’-CCAGACTTTTGCTACTGGGAC -3’ and RT-Itgal_1R 5’-GCTTGTTCGGCAGTGATAGAG – 3’) were designed using publicly available genomic sequences for C57BL/6J, WSB/EiJ and CC042 (48). Each reaction was performed in technical triplicates and Ct values were generated on a Viia7 Real-time PCR machine (Life Technologies). Relative expression for *Itgal* was calculated by using ΔΔCt for each sample relative to B-actin. Itgal PCR products were separated by gel electrophoresis on a 3% agarose gel and their sequences determined by Sanger sequencing.

## Statistical analysis

Statistical tests were performed using GraphPad Prism 7. Correlation between measured traits was visualized using corrplot v0.77 in R version 3.2.4 (Pearson correlation ordered by hclust).

## Supporting information

Table S1. Phenotype data for F2 intercross mice

Table S2. MiniMUGA Genotype data for CC001, CC042 and F2 intercross mice

## Data availability

All relevant data to support the findings of this study are located within the paper and supplemental files.

## Acknowledgements

We thank the Sassetti and Behar lab members for technical assistance, including Fred Boehm for statistical expertise; the UMASS Department of Animal Medicine for expert technical services and Christina Baer and the Sanderson Center for Optical Experimentation (SCOPE) imaging facility at UMass Medical School for assistance with microscopy. The TissueGnostics TissueFAXS PLUS slide scanning microscope was generously loaned to the SCOPE by TissueGnostics. This paper is dedicated to Riley Kay Proulx (MKPxSLP F_1_), whose gestation and arrival into the world coincided with the making, genotyping and analysis of this cross.

## Funding sources

This work was supported by a grant from the National Institutes of Health (AI132130) to C. Sassetti, F. Pardo-Manuel de Villena, S. Behar, and a fellowship from the Charles H. King Foundation to C. Smith.

---

**Table S1. Phenotype data for F_2_ intercross mice.** Smith_et_al_phenotypes.xls

**Table S2. MiniMUGA Genotype data for CC001, CC042 and F2 intercross mice.** Smith_et_al_genotypes.xls

## References

1. Houben RMGJ, Dodd PJ. 2016. The Global Burden of Latent Tuberculosis Infection: A Re-estimation Using Mathematical Modelling. PLoS Med 13:e1002152.

2. Comstock GW. 1978. Tuberculosis in twins: a re-analysis of the Prophit survey. Am Rev Respir Dis 117:621–624.

3. Kallmann FJ. 1943. Genetic mechanisms in resistance to tuberculosis. Psych Quar 17:32–37.

4. Bogunovic D, Byun M, Durfee LA, Abhyankar A, Sanal O, Mansouri D, Salem S, Radovanovic I, Grant AV, Adimi P, Mansouri N, Okada S, Bryant VL, Kong X-F, Kreins A, Velez MM, Boisson B, Khalilzadeh S, Ozcelik U, Darazam IA, Schoggins JW, Rice CM, Al-Muhsen S, Behr M, Vogt G, Puel A, Bustamante J, Gros P, Huibregtse JM, Abel L, Boisson-Dupuis S, Casanova J-L. 2012. Mycobacterial disease and impaired IFN-γ immunity in humans with inherited ISG15 deficiency. Science 337:1684–1688.

5. Bustamante J, Arias AA, Vogt G, Picard C, Galicia LB, Prando C, Grant AV, Marchal CC, Hubeau M, Chapgier A, de Beaucoudrey L, Puel A, Feinberg J, Valinetz E, Jannière L, Besse C, Boland A, Brisseau J-M, Blanche S, Lortholary O, Fieschi C, Emile J-F, Boisson-Dupuis S, Al-Muhsen S, Woda B, Newburger PE, Condino-Neto A, Dinauer MC, Abel L, Casanova J-L. 2011. Germline CYBB mutations that selectively affect macrophages in kindreds with X-linked predisposition to tuberculous mycobacterial disease. Nat Immunol 12:213–221.

6. Filipe-Santos O, Bustamante J, Chapgier A, Vogt G, de Beaucoudrey L, Feinberg J, Jouanguy E, Boisson-Dupuis S, Fieschi C, Picard C, Casanova J-L. 2006. Inborn errors of IL-12/23- and IFN-gamma-mediated immunity: molecular, cellular, and clinical features. Seminars in Immunology 18:347–361.

7. Hambleton S, Salem S, Bustamante J, Bigley V, Boisson-Dupuis S, Azevedo J, Fortin A, Haniffa M, Ceron-Gutierrez L, Bacon CM, Menon G, Trouillet C, McDonald D, Carey P, Ginhoux F, Alsina L, Zumwalt TJ, Kong X-F, Kumararatne D, Butler K, Hubeau M, Feinberg J, Al-Muhsen S, Cant A, Abel L, Chaussabel D, Doffinger R, Talesnik E, Grumach A, Duarte A, Abarca K, Moraes-Vasconcelos D, Burk D, Berghuis A, Geissmann F, Collin M, Casanova J-L, Gros P. 2011. IRF8 mutations and human dendritic-cell immunodeficiency. N Engl J Med 365:127–138.

8. Salem S, Gros P. 2013. Genetic determinants of susceptibility to Mycobacterial infections: IRF8, a new kid on the block. Adv Exp Med Biol 783:45–80.

9. Bellamy R, Ruwende C, Corrah T, McAdam KP, Whittle HC, Hill AV. 1998. Variations in the NRAMP1 gene and susceptibility to tuberculosis in West Africans. N Engl J Med 338:640–644.

10. Curtis J, Luo Y, Zenner HL, Cuchet-Lourenço D, Wu C, Lo K, Maes M, Alisaac A, Stebbings E, Liu JZ, Kopanitsa L, Ignatyeva O, Balabanova Y, Nikolayevskyy V, Baessmann I, Thye T, Meyer CG, Nürnberg P, Horstmann RD, Drobniewski F, Plagnol V, Barrett JC, Nejentsev S. 2015. Susceptibility to tuberculosis is associated with variants in the ASAP1 gene encoding a regulator of dendritic cell migration. Nat Genet 47:523–527.

11. Thye T, Owusu-Dabo E, Vannberg FO, van Crevel R, Curtis J, Sahiratmadja E, Balabanova Y, Ehmen C, Muntau B, Ruge G, Sievertsen J, Gyapong J, Nikolayevskyy V, Hill PC, Sirugo G, Drobniewski F, van de Vosse E, Newport M, Alisjahbana B, Nejentsev S, Ottenhoff THM, Hill AVS, Horstmann RD, Meyer CG. 2012. Common variants at 11p13 are associated with susceptibility to tuberculosis. Nat Genet 44:257–259.

12. Thye T, Vannberg FO, Wong SH, Owusu-Dabo E, Osei I, Gyapong J, Sirugo G, Sisay-Joof F, Enimil A, Chinbuah MA, Floyd S, Warndorff DK, Sichali L, Malema S, Crampin AC, Ngwira B, Teo YY, Small K, Rockett K, Kwiatkowski D, Fine PE, Hill PC, Newport M, Lienhardt C, Adegbola RA, Corrah T, Ziegler A, Morris AP, Meyer CG, Horstmann RD, Hill AVS. 2010. Genome-wide association analyses identifies a susceptibility locus for tuberculosis on chromosome 18q11.2. Nat Genet 42:739–741.

13. Medina E, North RJ. 1998. Resistance ranking of some common inbred mouse strains to Mycobacterium tuberculosis and relationship to major histocompatibility complex haplotype and Nramp1 genotype. Immunology 93:270–274.

14. Cooper AM, Dalton DK, Stewart TA, Griffin JP, Russell DG, Orme IM. 1993. Disseminated tuberculosis in interferon gamma gene-disrupted mice. J Exp Med 178:2243–2247.

15. Flynn JL, Chan J, Triebold KJ, Dalton DK, Stewart TA, Bloom BR. 1993. An essential role for interferon gamma in resistance to Mycobacterium tuberculosis infection. J Exp Med 178:2249–2254.

16. MacMicking JD, Taylor GA, McKinney JD. 2003. Immune control of tuberculosis by IFN-gamma-inducible LRG-47. Science 302:654–659.

17. Schaible UE, Sturgill-Koszycki S, Schlesinger PH, Russell DG. 1998. Cytokine activation leads to acidification and increases maturation of Mycobacterium avium-containing phagosomes in murine macrophages. The Journal of Immunology 160:1290–1296.

18. Desvignes L, Ernst JD. 2009. Interferon-gamma-responsive nonhematopoietic cells regulate the immune response to Mycobacterium tuberculosis. Immunity 31:974–985.

19. Nandi B, Behar SM. 2011. Regulation of neutrophils by interferon-γ limits lung inflammation during tuberculosis infection. J Exp Med 208:2251–2262.

20. Mishra BB, Lovewell RR, Olive AJ, Zhang G, Wang W, Eugenin E, Smith CM, Phuah JY, Long JE, Dubuke ML, Palace SG, Goguen JD, Baker RE, Nambi S, Mishra R, Booty MG, Baer CE, Shaffer SA, Dartois V, McCormick BA, Chen X, Sassetti CM. 2017. Nitric oxide prevents a pathogen-permissive granulocytic inflammation during tuberculosis. Nat Microbiol 2:17072.

21. Smith CM, Sassetti CM. 2018. Modeling Diversity: Do Homogeneous Laboratory Strains Limit Discovery? Trends in Microbiology 26:892–895.

22. Roy Chowdhury R, Vallania F, Yang Q, Lopez Angel CJ, Darboe F, Penn-Nicholson A, Rozot V, Nemes E, Malherbe ST, Ronacher K, Walzl G, Hanekom W, Davis MM, Winter J, Chen X, Scriba TJ, Khatri P, Chien Y-H. 2018. A multi-cohort study of the immune factors associated with M. tuberculosis infection outcomes. Nature 560:644–648.

23. Lu LL, Smith MT, Yu KKQ, Luedemann C, Suscovich TJ, Grace PS, Cain A, Yu WH, McKitrick TR, Lauffenburger D, Cummings RD, Mayanja-Kizza H, Hawn TR, Boom WH, Stein CM, Fortune SM, Seshadri C, Alter G. 2019. IFN-γ-independent immune markers of Mycobacterium tuberculosis exposure. Nature Medicine 25:977–987.

24. Lu LL, Chung AW, Rosebrock TR, Ghebremichael M, Yu WH, Grace PS, Schoen MK, Tafesse F, Martin C, Leung V, Mahan AE, Sips M, Kumar MP, Tedesco J, Robinson H, Tkachenko E, Draghi M, Freedberg KJ, Streeck H, Suscovich TJ, Lauffenburger DA, Restrepo BI, Day C, Fortune SM, Alter G. 2016. A Functional Role for Antibodies in Tuberculosis. Cell 167:433–443.e14.

25. Li H, Wang X-X, Wang B, Fu L, Liu G, Lu Y, Cao M, Huang H, Javid B. 2017. Latently and uninfected healthcare workers exposed to TB make protective antibodies against Mycobacterium tuberculosis. Proc Natl Acad Sci USA 114:5023–5028.

26. Chackerian AA, Behar SM. 2003. Susceptibility to Mycobacterium tuberculosis: lessons from inbred strains of mice. Tuberculosis 83:279–285.

27. Nadezhda Logunova MKVPAA. 2015. The QTL within the H2 Complex Involved in the Control of Tuberculosis Infection in Mice Is the Classical Class II H2-Ab1 Gene 1–22.

28. Kramnik I, Dietrich WF, Demant P, Bloom BR. 2000. Genetic control of resistance to experimental infection with virulent Mycobacterium tuberculosis. Proc Natl Acad Sci USA 97:8560–8565.

29. Pan H, Yan B-S, Rojas M, Shebzukhov YV, Zhou H, Kobzik L, Higgins DE, Daly MJ, Bloom BR, Kramnik I. 2005. Ipr1 gene mediates innate immunity to tuberculosis. Nature 434:767–772.

30. Sissons J, Yan B-S, Pichugin AV, Kirby A, Daly MJ, Kramnik I. 2008. Multigenic control of tuberculosis resistance: analysis of a QTL on mouse chromosome 7 and its synergism with sst1. Genes Immun 10:37–46.

31. Mitsos L-M, Cardon LR, Ryan L, LaCourse R, North RJ, Gros P. 2003. Susceptibility to tuberculosis: a locus on mouse chromosome 19 (Trl-4) regulates Mycobacterium tuberculosis replication in the lungs. Proc Natl Acad Sci USA 100:6610–6615.

32. Marquis JF, LaCourse R, Ryan L, North RJ, Gros P. 2009. Genetic and Functional Characterization of the Mouse Trl3 Locus in Defense against Tuberculosis. The Journal of Immunology 182:3757–3767.

33. Svenson KL, Gatti DM, Valdar W, Welsh CE, Cheng R, Chesler EJ, Palmer AA, McMillan L, Churchill GA. 2012. High-resolution genetic mapping using the Mouse Diversity outbred population. Genetics 190:437–447.

34. Niazi MKK, Dhulekar N, Schmidt D, Major S, Cooper R, Abeijon C, Gatti DM, Kramnik I, Yener B, Gurcan M, Beamer G. 2015. Lung necrosis and neutrophils reflect common pathways of susceptibility to Mycobacterium tuberculosis in genetically diverse, immune-competent mice. Dis Model Mech 8:1141–1153.

35. Churchill GA, Airey DC, Allayee H, Angel JM, Attie AD, Beatty J, Beavis WD, Belknap JK, Bennett B, Berrettini W, Bleich A, Bogue M, Broman KW, Buck KJ, Buckler E, Burmeister M, Chesler EJ, Cheverud JM, Clapcote S, Cook MN, Cox RD, Crabbe JC, Crusio WE, Darvasi A, Deschepper CF, Doerge RW, Farber CR, Forejt J, Gaile D, Garlow SJ, Geiger H, Gershenfeld H, Gordon T, Gu J, Gu W, de Haan G, Hayes NL, Heller C, Himmelbauer H, Hitzemann R, Hunter K, Hsu H-C, Iraqi FA, Ivandic B, Jacob HJ, Jansen RC, Jepsen KJ, Johnson DK, Johnson TE, Kempermann G, Kendziorski C, Kotb M, Kooy RF, Llamas B, Lammert F, Lassalle J-M, Lowenstein PR, Lu L, Lusis A, Manly KF, Marcucio R, Matthews D, Medrano JF, Miller DR, Mittleman G, Mock BA, Mogil JS, Montagutelli X, Morahan G, Morris DG, Mott R, Nadeau JH, Nagase H, Nowakowski RS, O’Hara BF, Osadchuk AV, Page GP, Paigen B, Paigen K, Palmer AA, Pan H-J, Peltonen-Palotie L, Peirce J, Pomp D, Pravenec M, Prows DR, Qi Z, Reeves RH, Roder J, Rosen GD, Schadt EE, Schalkwyk LC, Seltzer Z, Shimomura K, Shou S, Sillanpää MJ, Siracusa LD, Snoeck H-W, Spearow JL, Svenson K, Tarantino LM, Threadgill D, Toth LA, Valdar W, de Villena FP-M, Warden C, Whatley S, Williams RW, Wiltshire T, Yi N, Zhang D, Zhang M, Zou F, Complex Trait Consortium. 2004. The Collaborative Cross, a community resource for the genetic analysis of complex traits. Nat Genet 36:1133–1137.

36. Collaborative Cross Consortium. 2012. The genome architecture of the Collaborative Cross mouse genetic reference population. Genetics 190:389–401.

37. Smith CM, Proulx MK, Olive AJ, Laddy D, Mishra BB, Moss C, Gutierrez NM, Bellerose MM, Barreira-Silva P, Phuah JY, Baker RE, Behar SM, Kornfeld H, Evans TG, Beamer G, Sassetti CM. 2016. Tuberculosis Susceptibility and Vaccine Protection Are Independently Controlled by Host Genotype. mBio 7:e01516–16.

38. Sakai S, Kauffman KD, Sallin MA, Sharpe AH, Young HA, Ganusov VV, Barber DL. 2016. CD4 T Cell-Derived IFN-γ Plays a Minimal Role in Control of Pulmonary Mycobacterium tuberculosis Infection and Must Be Actively Repressed by PD-1 to Prevent Lethal Disease. PLoS Pathog 12:e1005667.

39. Shorter JR, Najarian ML, Bell TA, Blanchard M, Ferris MT, Hock P, Kashfeen A, Kirchoff KE, Linnertz CL, Sigmon JS, Miller DR, McMillan L, Pardo-Manuel de Villena F. 2019. Whole Genome Sequencing and Progress Toward Full Inbreeding of the Mouse Collaborative Cross Population. G3&amp;#58; Genes|Genomes|Genetics 9:1303–1311.

40. Aylor DL, Valdar W, Foulds-Mathes W, Buus RJ, Verdugo RA, Baric RS, Ferris MT, Frelinger JA, Heise M, Frieman MB, Gralinski LE, Bell TA, Didion JD, Hua K, Nehrenberg DL, Powell CL, Steigerwalt J, Xie Y, Kelada SNP, Collins FS, Yang IV, Schwartz DA, Branstetter LA, Chesler EJ, Miller DR, Spence J, Liu EY, McMillan L, Sarkar A, Wang J, Wang W, Zhang Q, Broman KW, Korstanje R, Durrant C, Mott R, Iraqi FA, Pomp D, Threadgill D, de Villena FP-M, Churchill GA. 2011. Genetic analysis of complex traits in the emerging Collaborative Cross. Genome Research 21:1213–1222.

41. Baaten BJG, Li C-R, Deiro MF, Lin MM, Linton PJ, Bradley LM. 2010. CD44 regulates survival and memory development in Th1 cells. Immunity 32:104–115.

42. Seder RA, Darrah PA, Roederer M. 2008. T-cell quality in memory and protection: implications for vaccine design. Nat Rev Immunol 8:247–258.

43. Szafranski K, Schindler S, Taudien S, Hiller M, Huse K, Jahn N, Schreiber S, Backofen R, Platzer M. 2007. Violating the splicing rules: TG dinucleotides function as alternative 3’ splice sites in U2-dependent introns. Genome Biol 8:R154–11.

44. Ghosh S, Chackerian AA, Parker CM, Ballantyne CM, Behar SM. 2006. The LFA-1 adhesion molecule is required for protective immunity during pulmonary Mycobacterium tuberculosis infection. The Journal of Immunology 176:4914–4922.

45. Sallin MA, Kauffman KD, Riou C, Bruyn Du E, Foreman TW, Sakai S, Hoft SG, Myers TG, Gardina PJ, Sher A, Moore R, Wilder-Kofie T, Moore IN, Sette A, Lindestam Arlehamn CS, Wilkinson RJ, Barber DL. 2018. Host resistance to pulmonary Mycobacterium tuberculosis infection requires CD153 expression. Nat Microbiol 3:1198–1205.

46. Gopal R, Monin L, Slight S, Uche U, Blanchard E, A Fallert Junecko B, Ramos-Payan R, Stallings CL, Reinhart TA, Kolls JK, Kaushal D, Nagarajan U, Rangel-Moreno J, Khader SA. 2014. Unexpected Role for IL-17 in Protective Immunity against Hypervirulent Mycobacterium tuberculosis HN878 Infection. PLoS Pathog 10:e1004099–14.

47. Threadgill DW, Churchill GA. 2012. Ten years of the Collaborative Cross. Genetics 190:291–294.

48. Srivastava A, Morgan AP, Najarian ML, Sarsani VK, Sigmon JS, Shorter JR, Kashfeen A, McMullan RC, Williams LH, Giusti-Rodríguez P, Ferris MT, Sullivan P, Hock P, Miller DR, Bell TA, McMillan L, Churchill GA, de Villena FP-M. 2017. Genomes of the Mouse Collaborative Cross. Genetics 206:537–556.

49. Welsh CE, Miller DR, Manly KF, Wang J, McMillan L, Morahan G, Mott R, Iraqi FA, Threadgill DW, de Villena FP-M. 2012. Status and access to the Collaborative Cross population. Mamm Genome 23:706–712.

