## Supplemental Figures 1-3 for "Functionally overlapping variants control TB susceptibility in Collaborative Cross mice"

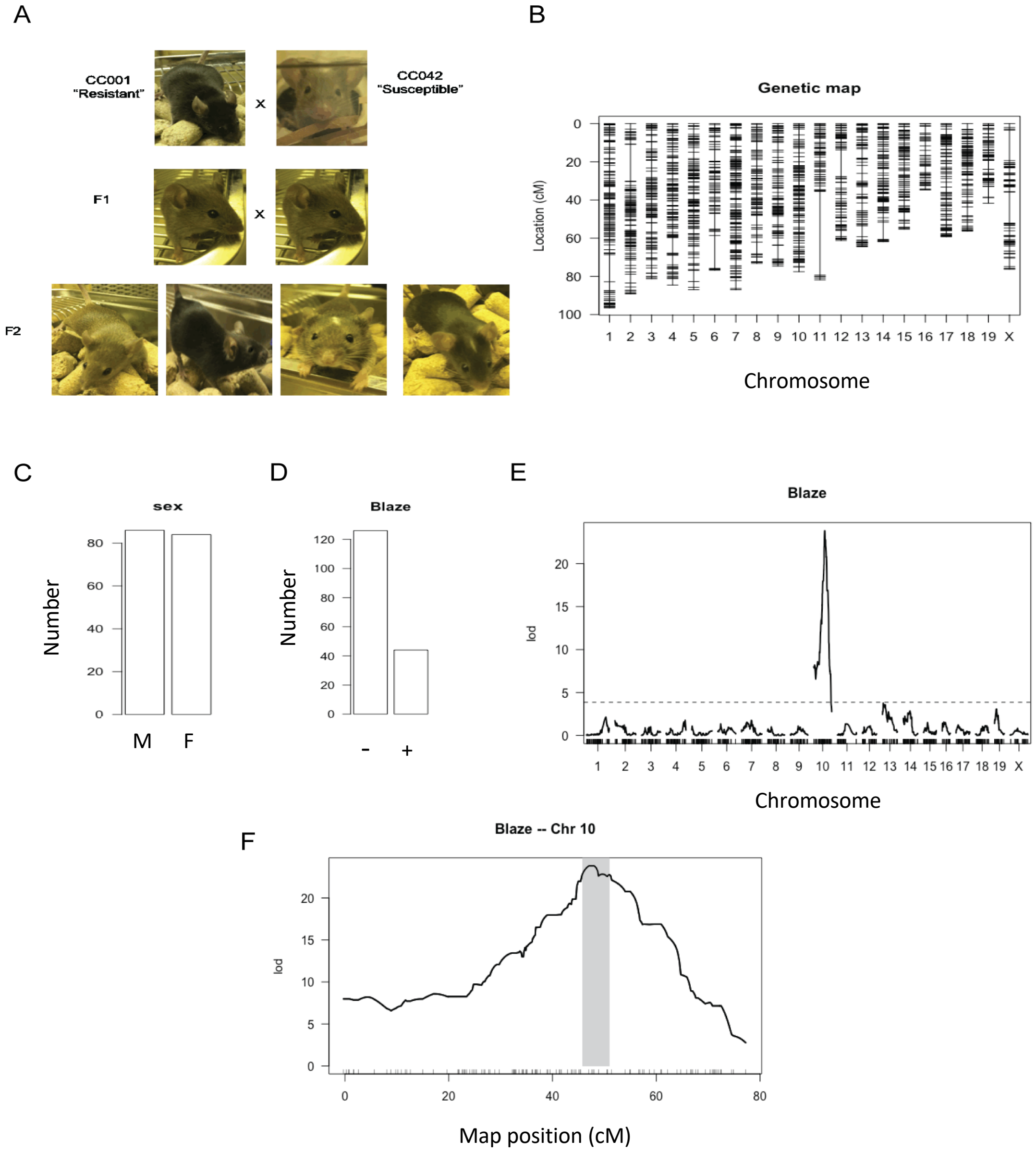


**Supplemental Figure 1. Description of F2 cross population and test scan for the Mendelian "Blaze" trait.** (A) Coat color phenotypes of parent mice, CC001 (black), CC042 (agouti with white head blaze), $F_1$ (agouti) and $F_2$ (agouti, black, agouti and head blaze, black and head blaze). (B) genetic map of F2 population used in QTL studies. Vertical lines show Chromosomes (1-19 autosomes, X = X Chromosome). Horizontal ticks show marker location in cM. (C) Number of male vs. female mice (male=M, female=F). (D) Number of mice with no head blaze vs head blaze in F2 mice (no head blaze= - ; head blaze= +). (E) Results of genome scan for "Blaze" trait. (F) Bayes interval for the "Blaze" trait on chromosome 7 (shaded) containing *Kitl*, previously shown to be associated with the $WSB^{blaze}$ phenotype (Aylor DL *et al.* 2011. Genome Research 21:1213–1222).

A

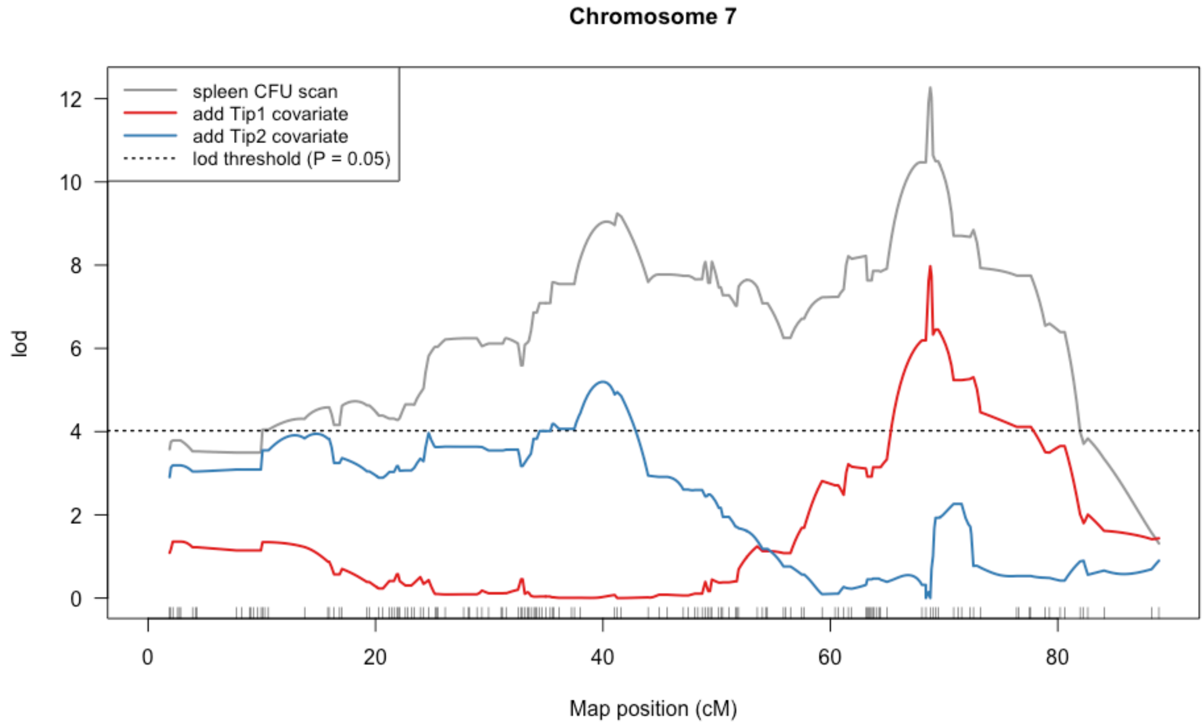


B

| | df | Type III SS | LOD | %var | F value | Pvalue(Chi2) | Pvalue(F) |
|---|---|---|---|---|---|---|---|
| Tip1 | 2 | 5.78 | 4.99 | 6.79 | 11.52 | 1.01E-05 | 2.14E-05 |
| Tip2 | 2 | 10.37 | 8.53 | 12.19 | 20.67 | 2.95E-09 | 1.06E-08 |
| Tip3 | 2 | 2.55 | 2.29 | 2.99 | 5.08 | 5.18E-03 | 7.30E-03 |

**Supplemental Figure 2. Tests for independence of Tip1 and Tip2 QTL underlying spleen CFU.** (A) The spleen CFU trait was re-mapped using the genotype probabilities at Tip1 and Tip2 separately as covariates. The distal QTL (Tip2) QTL reaches significant LOD after variation explained by Tip1 was removed and vice versa. (B) A multi-QTL model including Tip1, Tip2, and Tip3 was fit for the spleen CFU phenotype (batch and sex included as covariates). The table shows results from drop-one-term ANOVA where each QTL is dropped from the model, one at a time, and the sub-model with that factor omitted is compared to the full model. The results provide substantial evidence for all QTL. Column headings: df, degrees of freedom; Type III SS, Type III sum of squares; LOD, LOD score; %var, percentage of variance explained; Pvalue(Chi2), P value for chi square; Pvalue(F), P value for F distribution. The degree of freedom, Type III sum of squares, LOD score, and percentage of variance explained are the values comparing the full to the sub-model.

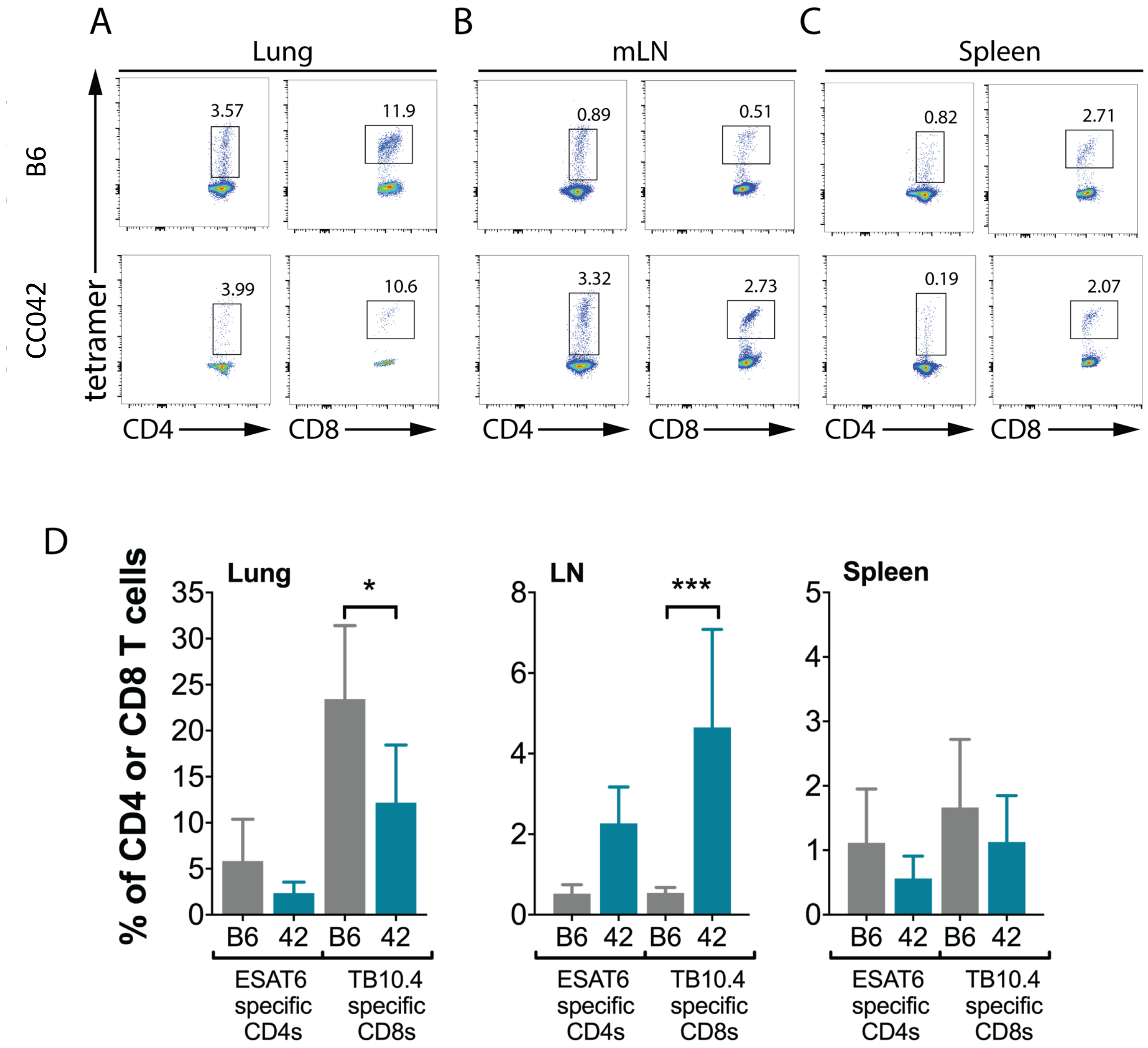


**Supplemental Figure 3. Frequency of antigen-specific T cells in B6 and CC042 following pulmonary Mtb infection**. Representative flow plots showing the frequencies of ESAT-6-specific CD4 T cells and TB10.4-specific CD8 T cells in the (A) Lung (B) mediastinal lymph node (mLN) and (C) spleen of B6 (grey shading) and CC042 (teal shading) mice at 4 weeks post pulmonary Mtb infection. Bar plots show mean + SD of ESAT-6-specific CD4 T cells and TB10.4-specific CD8 T cells in the (D) Lung (E) mLN and (F) spleen at the same timepoint. Sidak's multiple comparison test was used to determine significance where $p<0.05$ *, $p<0.001$***.
